# Investigating the relationship between insecticide resistance, underlying molecular mechanisms and malaria prevalence in *Anopheles gambiae* s.l. from Guinea

**DOI:** 10.1101/434688

**Authors:** Emma Collins, Natasha M. Vaselli, Moussa Sylla, Abdoul H. Beavogui, James Orsborne, Thomas Walker, Louisa A. Messenger

## Abstract

The threat of insecticide resistance across sub-Saharan Africa is anticipated to severely impact the continued effectiveness of malaria vector control. We investigated the effect of carbamate and pyrethroid resistance on *Anopheles gambiae* s.l age, *Plasmodium falciparum* infection and characterized molecular resistance mechanisms in Guinea. Pyrethroid resistance was intense, with survivors of ten times the insecticidal concentration required to kill susceptible individuals. The L1014F *kdr* allele was significantly associated with mosquito survival following deltamethrin or permethrin treatment (*p*=0.003 and *p*=0.04, respectively). N1575Y and I1527T mutations were identified in 13% and 10% of individuals, respectively, but neither conferred increased pyrethroid tolerance. Partial restoration of pyrethroid susceptibility following synergist pre-exposure suggest a role for mixed-function oxidases. Carbamate resistance was lower and significantly associated with the G119S *Ace-1* mutation (*p*=0.001). Oocyst rates were 6.8% and 4.2% among resistant and susceptible mosquitoes, respectively; survivors of bendiocarb exposure were significantly more likely to be infected (*p*=0.03). Resistant mosquitoes had significantly lower parity rates; however, a subset of intensely pyrethroid-resistant vectors were more likely to be parous (*p*=0.042 and *p*=0.045, for survivors of five and ten times the diagnostic dose of insecticides, respectively). Our findings emphasize the need for additional studies directly assessing the influence of insecticide resistance on mosquito fitness.

## Introduction

Malaria remains a leading cause of morbidity and mortality in the tropics, where it is estimated to have resulted in ~445,000 deaths in 2016 alone^1^. Despite considerable reductions in disease burden achieved by scaling-up the provision of long-lasting insecticidal nets (LLINs) and indoor residual spraying (IRS)^2^, future, long-term effectiveness of both strategies may be jeopardized by widespread emergence of insecticide resistance throughout mosquito populations^3,4^. In response, there is a growing impetus among commercial manufacturers to develop alternate vector control interventions, including novel insecticide classes, combinations and formulations, as well as National Malaria Control Programmes (NMCPs) and international policy makers to expand and heighten resistance monitoring and surveillance^5^. However, the severity of this threat is currently unknown because there is limited evidence linking the operational failure of available control measures to the presence of local, resistant mosquito species^6,7,8,9,10,11,12,13^.

The efficacy of IRS and LLINs is predicated on their ability to reduce the daily survival rate of *Anopheles* mosquitoes and prevent the completion of parasite development to the infectious stage. In the context of LLINs, one potential explanation for their continued effectiveness in malaria endemic regions, is that intact nets provide a physical barrier to mosquito feeding, even in the presence of increased vector tolerance to their insecticidal properties. A meta-analysis of field data indicate that treated nets reduced blood feeding and increased mosquito mortality compared to untreated nets, even in areas with the highest levels of resistance^4^. Furthermore, a recent large-scale, multi-country trial reported no association between malaria disease burden and pyrethroid resistance, with evidence that LLINs continued to provide personal protection across areas of different resistance intensities^14^. Laboratory studies now suggest that pleiotropic fitness costs associated with insecticide resistance may influence malaria transmission either by directly reducing mosquito life span^15^ and/or fecundity^16^, altering host seeking, feeding and mating behaviours^17 18 19^ or by impairing parasite development inside vectors^20 21 22^. However, to date, few field studies have directly investigated the impact of insecticide resistance intensity on malaria transmission dynamics.

In Guinea, malaria represents a significant public health problem, where the entire population of ~11.7 million people is at risk and the nationwide prevalence is estimated at 15%^23^. Between 2013-2017, the NMCP, with support from the President’s Malaria Initiative (PMI) and the Global Fund, have procured and distributed over 27.6 million pyrethroid LLINs^24^. Because the current vector control strategy relies almost exclusively on LLIN use^25^, nationwide pyrethroid resistance, is of significant concern^26^. To better inform future malaria control efforts in Guinea, there is a pressing need to characterize levels of operationally-significant insecticide resistance, as well as determine the effect this phenomenon has on the vectorial capacity of local mosquitoes to transmit malaria.

## Results

### Mosquito abundance and species identification

A total of 3962 female *An. gambiae* s.l. mosquitoes were captured from six sites in Maferinyah subprefecture, in Forecariah Prefecture (Senguelen=766; Yindi=755; Maferinyah Centre I=660; Madinagbe=608; Fandie=608 and Moribayah=565) over 25 days by manual aspiration from houses and human landing catches. A subsample of 181 were selected for molecular form identification, of which 68% (123/181) and 31% (56/181) were determined to be *An. coluzzii* (M molecular form) and *An. gambiae s.s.* (S molecular form), respectively; two hybrid forms (*An. gambiae*-*An. coluzzii*) were also identified. While both molecular forms were sympatric across Forecariah Prefecture, proportions did differ significantly among study villages (χ^2^=26.99; *p*=0.003), potentially attributable to varying local ecological factors favoured by each form^27^(Figure 1A).

**Figure 1.**
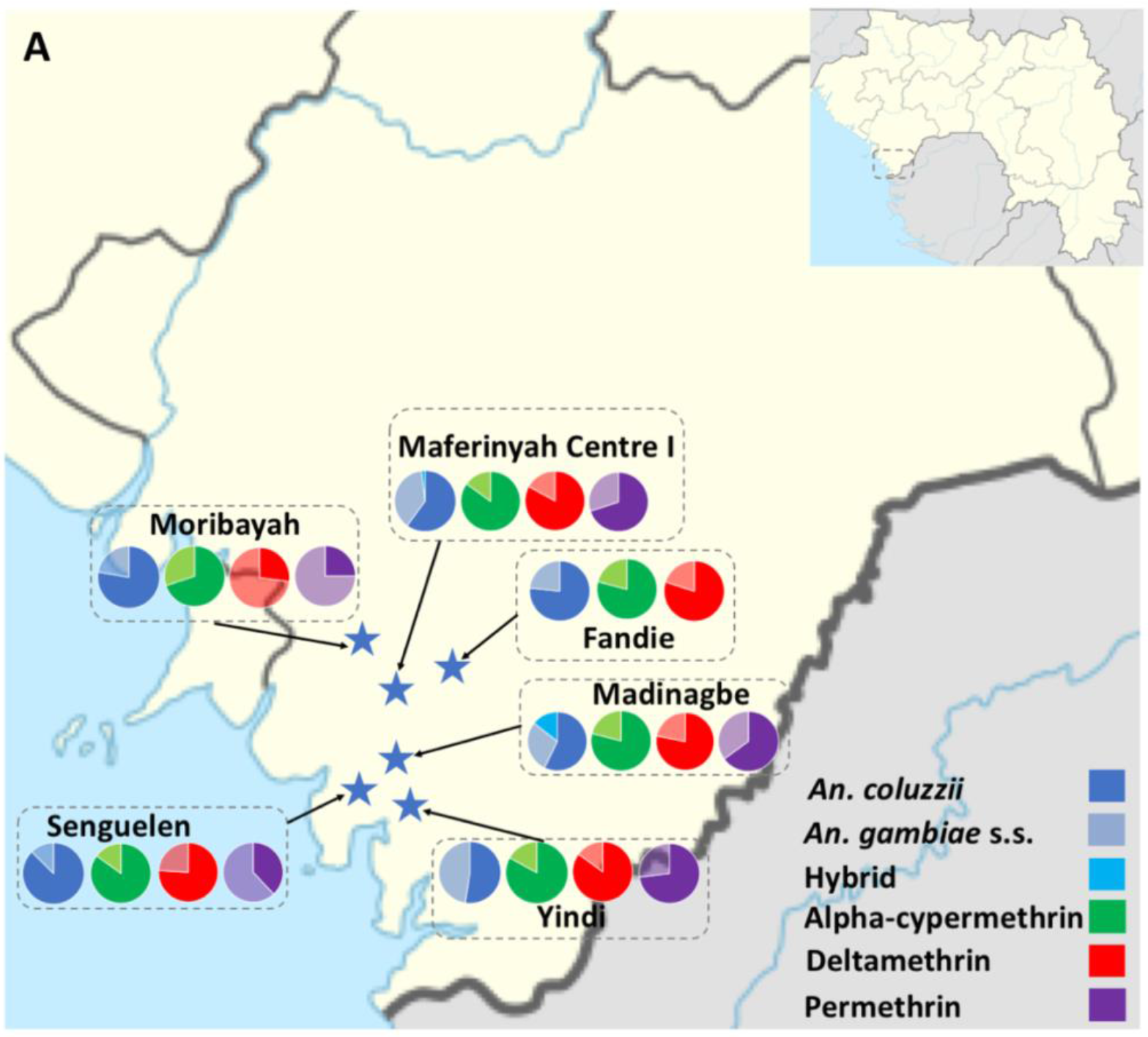
**A:**Map of Forecariah Prefecture, Guinea displaying proportions of *An. Gambiae* s.l. molecular forms and susceptibility levels to the diagnostic dose (1X) of alpha-cypermethrin, deltamethrin and permethrin, measured using CDC bottle bioassays, at six study sites indicated by stars (Fandie, Madinagbe, Maferinyah Centre I, Moribayah, Senguelen and Yindi). Inset map shows the location of Forecariah Prefecture in Guinea. **B:**Map of Forecariah Prefecture, Guinea displaying frequencies of L1014F, N1575Y and I1527T resistant (R) and wild type (S) alleles. **C:**Map of Forecariah Prefecture, Guinea displaying susceptibility levels to the diagnostic dose (1X) of bendiocarb, measured using CDC bottle bioassays, and frequency of G119S *Ace-1* resistant (R) and wild type (S) alleles. For all maps, legend colours referring to insecticides have darker shading denoting average mosquito mortality.

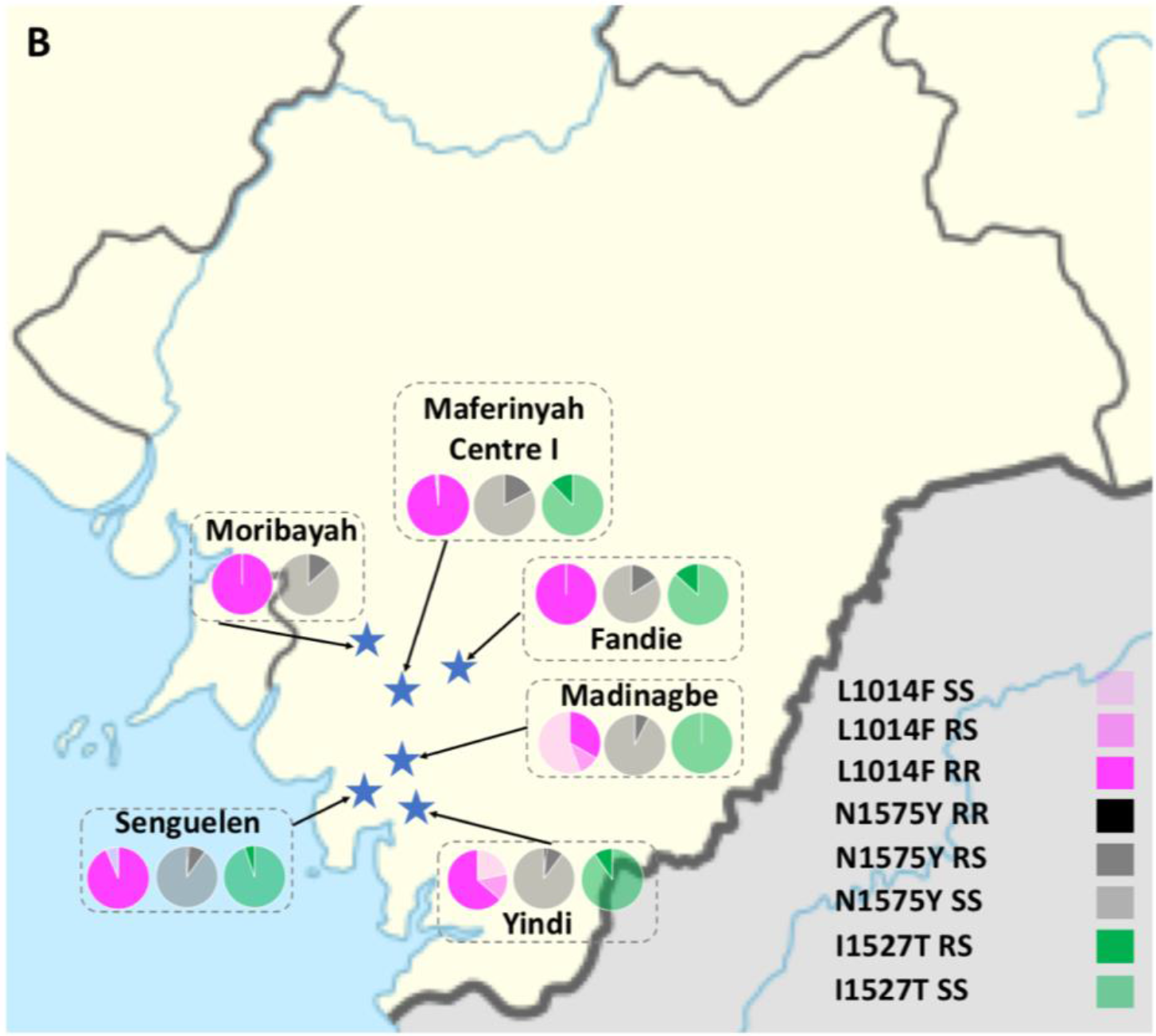

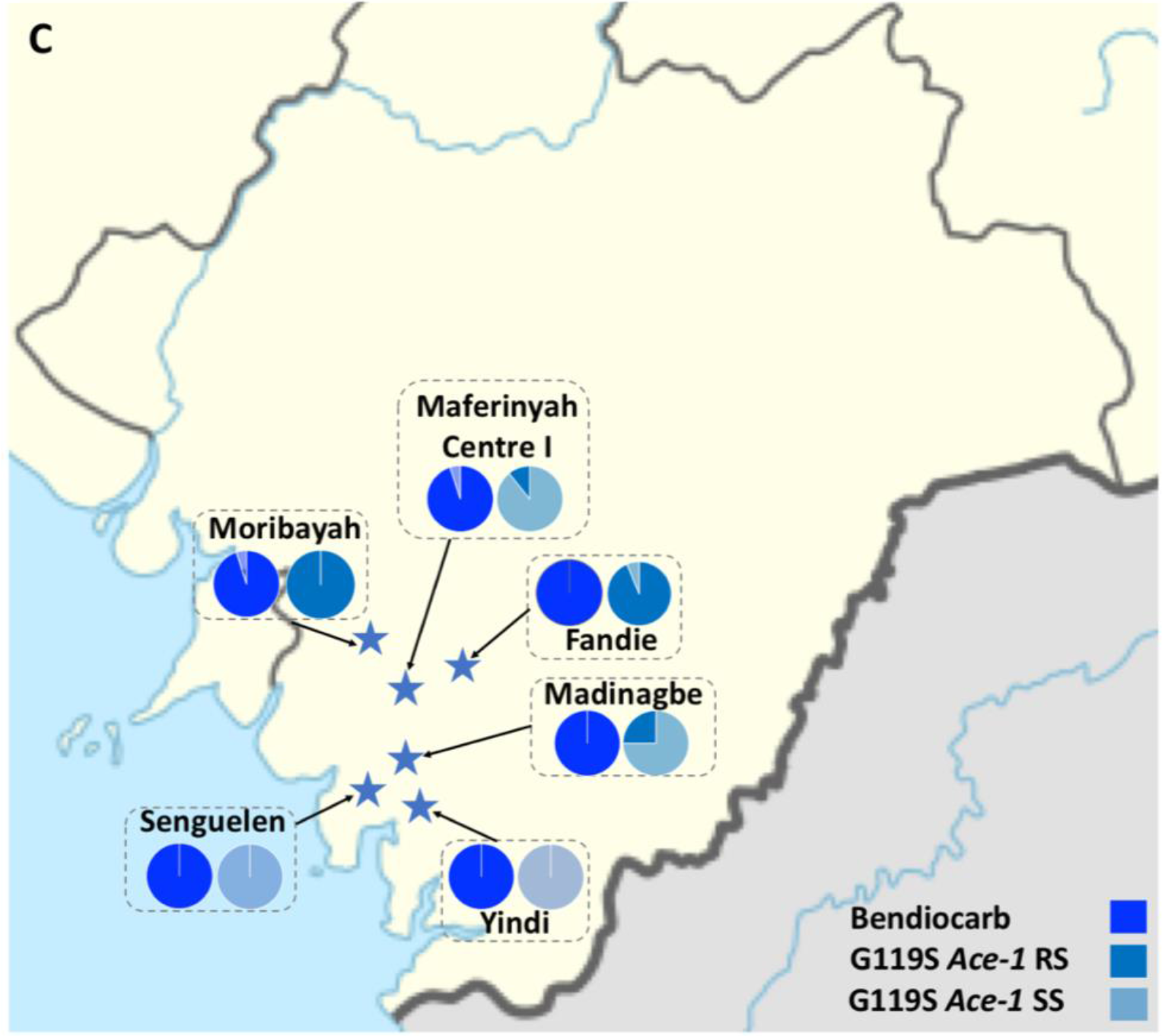

### Insecticide resistance intensity

Levels of resistance to four insecticides (alpha-cypermethrin, bendiocarb, deltamethrin and permethrin) were assessed among 2229 female *An. gambiae* s.l. mosquitoes, collected across six study sites in Forecariah Prefecture (Figure 1A and Figure 1C). Local vectors were characterized by intense but highly variable pyrethroid resistance, with all populations demonstrating less than 90% mosquito mortality to the diagnostic doses of pyrethroids and most areas containing individuals capable of surviving exposure to ten times these insecticide concentrations (Table 1 and Figure 2). In addition, a subsample of mosquitoes still living following 30 minutes of insecticide exposure were held for up to two hours in treated bottles, with more than 30% of vectors capable of surviving this extended exposure time in Madinagbe (17/55) Moribayah (19/38), Seneguelen (30/82) and Yindi (9/27); there was no significant difference in survival at two hours between the three pyrethroids under evaluation (χ^2^=4.35; *p*=0.114). A significant decline in proportions of mosquitoes surviving two hours of pyrethroid exposure was observed with increasing insecticide concentration; 37% (38/102), 25% (24/95), 26% (21/81) and 10% (5/52) of individuals survived contact with 1X, 2X, 5X and 10X for two hours (χ^2^=13.70; *p*=0.003), respectively. The highest levels of pyrethroid resistance were observed in Moribayah, where mosquito mortality to two times the diagnostic doses of deltamethrin and permethrin was 38% and 32%, respectively (Table 1). In general, levels of resistance were greater to permethrin compared to alpha-cypermethrin or deltamethrin, e.g. in Senguelen mosquito mortality was 38% *vs.* 85% and 76%, respectively.

By comparison, levels of carbamate resistance were low, with complete mosquito susceptibility observed in Fandie, Madinagbe, Senguelen and Yindi; possible resistance (95% mosquito mortality) was restricted to two adjacent villages (Maferinyah Centre I and Moribayah) (Figure 1C and Table 1).

For all insecticides, there was no significant difference in ability to survive exposure between *An. coluzzii* and *An. gambiae* s.s. (χ^2^=3.60; *p*=0.165) or association with survival and physiological status (i.e. blood-fed, unfed, gravid, etc.) (χ^2^=0.76; *p*=0.999).

**Table 1.** Percentage corrected mortality (and numbers tested) of *Anopheles gambiae* s.l. in CDC resistance intensity and synergist assays conducted in six sites in Forecariah Prefecture, Guinea.

| Study Site | Insecticides | % Mortality (numbers tested) after the diagnostic time |  |  |  |
| --- | --- | --- | --- | --- | --- |
|  |  | 1X | 2X | 5X | 10X |
| Fandie | Alpha-cypermethrin | 79 (19) | 86 (21) | 83 (23) | 89 (19) |
|  | Alpha-cypermethrin + PBO | 91 (22) | 95 (19) | 95 (22) | 100 (13) |
|  | Bendiocarb | 100 (23) | - | - | - |
|  | Deltamethrin | 80 (20) | 78 (22) | 74 (19) | 95 (20) |
| Madinagbe | Alpha-cypermethrin | 79 (24) | 80 (25) | 88 (24) | 88 (17) |
|  | Bendiocarb | 100 (19) | - | - | - |
|  | Deltamethrin | 78 (23) | 89 (19) | 89 (18) | 86 (21) |
|  | Permethrin | 65 (20) | 71 (21) | 78 (27) | 78 (18) |
| Maferinyah Centre I | Alpha-cypermethrin | 85 (41) | 98 (43) | 94 (46) | 87 (38) |
|  | Bendiocarb | 95 (40) | - | - | - |
|  | Deltamethrin | 83 (35) | 89 (35) | 83 (35) | 97 (35) |
|  | Permethrin | 70 (20) | 95 (20) | 100 (21) | 95 (20) |
| Moribayah | Alpha-cypermethrin | 70 (20) | 86 (21) | 82 (22) | 87 (23) |
|  | Bendiocarb | 95 (20) | - | - | - |
|  | Deltamethrin | 27 (22) | 38 (21) | 86 (22) | 88 (16) |
|  | Permethrin | 25 (24) | 32 (25) | 86 (22) | 95 (20) |
| Seneguelen | Alpha-cypermethrin | 85 (41) | 88 (41) | 92 (37) | 93 (43) |
|  | Bendiocarb | 100 (22) | - | - | - |
|  | Deltamethrin | 76 (42) | 77 (44) | 86 (42) | 83 (41) |
|  | Deltamethrin + PBO | 81 (16) | 88 (16) | 89 (19) | 100 (11) |
|  | Permethrin | 38 (29) | 40 (25) | 96 (25) | 100 (19) |
| Yindi | Alpha-cypermethrin | 83 (42) | 95 (43) | 90 (30) | 92 (25) |
|  | Bendiocarb | 100 (22) | - | - | - |
|  | Deltamethrin | 85 (40) | 90 (39) | 92 (36) | 95 (42) |
|  | Permethrin | 73 (45) | 89 (36) | 92 (36) | 100 (35) |

**Figure 2.**
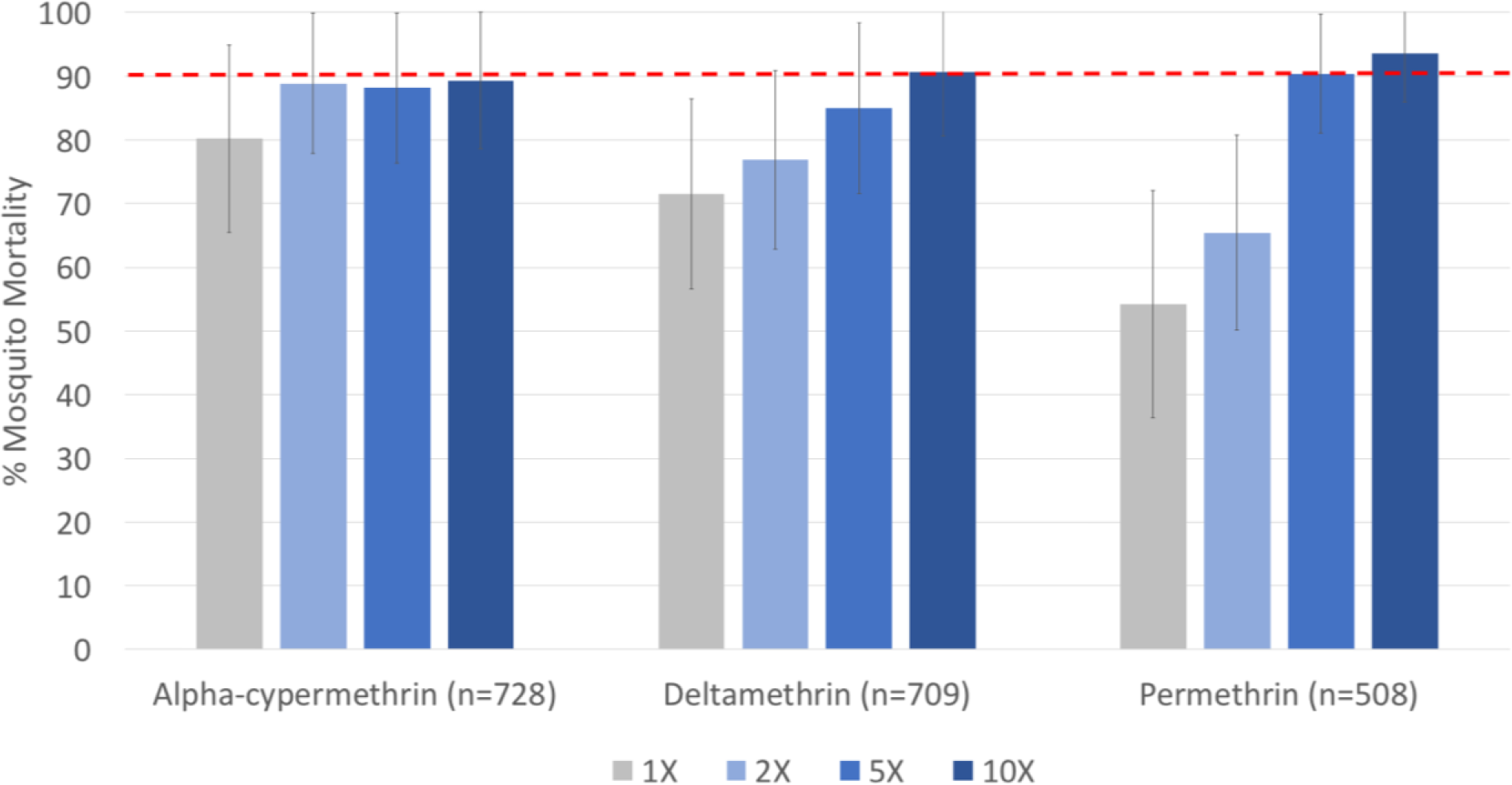
CDC resistance intensity assay data for three pyrethroid insecticides (alpha-cypermethrin, deltamethrin and permethrin) pooled across six study sites in Forecariah Prefecture (Fandie, Madinagbe, Maferinyah Centre I, Moribayah, Senguelen and Yindi). Mortality below 90% (indicated by the dashed red line) indicates the presence of confirmed resistance.

### Mosquito parity

Of 737 *An. gambiae* s.l. mosquitoes tested in resistance bioassays, which had their ovaries inspected for parity, 46% were nulliparous (340/737), i.e. had not laid an egg batch, and 54% (397/737) were parous. There was no significant difference in proportion of parous mosquitoes by study village (χ^2^=2.70; *p*=0.75). However, for all three pyrethroids under evaluation, resistant mosquitoes were significantly more likely to be nulliparous (alpha-cypermethrin: χ^2^=15.59; *p*<0.0000; deltamethrin: χ^2^=8.05; *p*=0.005; and permethrin: χ^2^=8.69; *p*=0.003). Small sample numbers precluded a direct comparison between pyrethroid resistance intensity and parity per insecticide, but when considering resistance levels of all pyrethroids, mosquitoes which were resistant to lower insecticide concentrations (2X) were significantly more likely to be nulliparous (χ^2^=17.78; *p*<0.0001), while greater proportions of those surviving exposure to five and ten times the diagnostic doses were parous (χ^2^=4.13; *p*=0.042 and χ^2^=4.03; *p*=0.045, respectively). Similarly, survivors of exposure to the diagnostic dose (1X) of bendiocarb were significantly more likely to be nulliparous (χ^2^=3.75; *p*=0.05).

### Target site and metabolic mechanisms of resistance

N1575Y mutation screening was undertaken in a subset of 388 *An. gambiae* s.l. and was detected in 13% (49/388) of samples in both *An. coluzzii* and *An. gambiae* s.s.; only two homozygote individuals were detected (Figure 1B and Table 2). The frequency of the N1575Y resistant allele varied non-significantly both among study districts, from 0.04 in Madinagbe to 0.09 in Maferinyah Centre I (χ^2^=8.34; *p*=0.60) (Table 2 and Figure 1B), and between mosquitoes surviving and dying in pyrethroid bioassays, respectively (χ^2^=2.05; *p*=0.36). There was no significant association between N1575Y allele frequency and ability of mosquitoes to survive pyrethroid exposure (alpha-cypermethrin: χ^2^=1.58; *p*=0.45; deltamethrin: χ^2^=0.79; *p*=0.375 and permethrin: χ^2^=1.52; *p*=0.22) or extended two hour pyrethroid exposure (χ^2^=2.47; *p*=0.29).

One hundred and twenty-eight base pairs upstream of N1575Y, a second non-synonymous mutation (I1527T) was identified among 10% of individuals (11/109); this mutation was not linked to N1575Y and only one individual from Maferinyah Centre I had both resistance mutations. All individuals with the I1527T resistant allele were heterozygous for this mutation and were identified as *An. gambiae* s.s. The frequency of the L1527T resistant allele varied non-significantly among study villages (χ^2^=1.48; *p*=0.83), ranging from 0 in Madinagbe to 0.07 in Fandie (Table 3 and Figure 1B). There was no significant association between I1527T allele frequency and ability of mosquitoes to survive pyrethroid exposure (alpha-cypermethrin: χ^2^=0.04; *p*=0.84; deltamethrin: χ^2^=0.22; *p*=0.64 and permethrin: χ^2^=2.94; *p*=0.09) or extended two hour pyrethroid exposure (χ^2^=0.13; *p*=0.72).

The presence of L1014F *kdr* allele was identified among 87% of samples (211/242), with the majority of mosquitoes presenting homozygous *kdr* profiles (83%; 200/242). The frequency of the L1014F *kdr* resistant allele varied significantly among study villages (χ^2^=93.82; *p*<0.0001), ranging from 0.39 in Madinagbe to 1.0 in Fandie and Moribayah (Table 4 and Figure 1B). There was a significant association between L1014F *kdr* allele frequency and ability of mosquitoes to survive deltamethrin or permethrin exposure (χ^2^=11.72; *p*=0.003 and χ^2^=6.39; *p*=0.04, respectively); however, this did not extend to survival after two hours of either insecticide treatment (χ^2^=1.45; *p*=0.48 and χ^2^=3.87; *p*=0.14, respectively). No significant association with the L1014F *kdr* allele frequency and ability to survive alpha-cypermethrin exposure was observed (χ^2^=1.67; *p*=0.43).

The presence of G119S *Ace-1* mutation was identified among 10% of samples (7/67) (Figure 1C). All individuals with the resistant allele were heterozygous for this mutation and were identified as *An. gambiae* s.s. Frequencies of the G119S *Ace-1* mutation varied significantly among study districts ranging from 0 in Senguelen and Yindi to 0.5 in Moribayah (χ^2^=20.34; *p*=0.001) (Table 5 and Figure 1C). There was a significant association between presence of the G119S *Ace-1* mutation and ability of mosquitoes to survive 30 minute and two hour carbamate exposure (χ^2^=11.51; *p*=0.001 and χ^2^=18.63; *p*<0.0001, respectively).

Across Forecariah Prefecture, significant deviations from Hardy-Weinberg equilibrium were observed for L1014F *kdr* in Madinagbe, Maferinyah Centre I, Senguelen and Yindi (*p*<0.0001 for all) (Table 4) but not for any other resistance loci.

Finally, to confirm the potential role of cytochrome P450 enzymes in pyrethroid resistance, mosquitoes collected in Fandie and Senguelen were pre-exposed to piperonyl butoxide (PBO), prior to intensity testing with alpha-cypermethrin and deltamethrin, respectively. In both populations, susceptibility was partially restored at five times the diagnostic dose of pyrethroids (95% and 89% mosquito mortality to alpha-cypermethrin and deltamethrin, respectively) and fully restored at 10X (Table 1).

**Table 2.** N1575Y allele frequencies in *An. gambiae* s.l. from six study sites in Forecariah Prefecture, Guinea.

| Study Site | # Mosquitoes tested | Homozygote mutation (RR) | Heterozygote mutation (RS) | Homozygote wild type (SS) | N1575Y allele frequency | | $\chi^2$ Test | P value |
| --- | --- | --- | --- | --- | --- | --- | --- | --- |
|  |  |  |  |  | R | S |  |  |
| Fandie | 63 | 0 | 10 | 53 | 0.08 | 0.92 | 0.47 | 0.493 |
| Madinagbe | 40 | 0 | 3 | 37 | 0.04 | 0.96 | 0.06 | 0.807 |
| Maferinyah Centre I | 87 | 0 | 15 | 72 | 0.09 | 0.91 | 0.77 | 0.380 |
| Moribayah | 38 | 0 | 5 | 33 | 0.07 | 0.93 | 0.19 | 0.663 |
| Seneguelen | 91 | 1 | 8 | 82 | 0.05 | 0.05 | 2.14 | 0.144 |
| Yindi | 69 | 1 | 6 | 62 | 0.06 | 0.94 | 2.87 | 0.090 |

**Table 3.** I1527T allele frequencies in *An. gambiae* s.l. from five study sites in Forecariah Prefecture, Guinea.

| Study Site | # Mosquitoes tested | Homozygote mutation (RR) | Heterozygote mutation (RS) | Homozygote wild type (SS) | I1527T allele frequency | | $\chi^2$ Test | P value |
| --- | --- | --- | --- | --- | --- | --- | --- | --- |
|  |  |  |  |  | R | S |  |  |
| Fandie | 15 | 0 | 2 | 13 | 0.07 | 0.93 | 0.08 | 0.777 |
| Madinagbe | 5 | 0 | 0 | 5 | 0 | 1.0 | - | - |
| Maferinyah Centre I | 49 | 0 | 6 | 43 | 0.06 | 0.94 | 0.21 | 0.647 |
| Seneguelen | 19 | 0 | 1 | 18 | 0.03 | 0.97 | 0.01 | 0.920 |
| Yindi | 21 | 0 | 2 | 19 | 0.05 | 0.95 | 0.05 | 0.823 |

**Table 4.** L1014F *kdr* allele frequencies in *An. gambiae* s.l. from six study sites in Forecariah Prefecture, Guinea.

| Study Site | # Mosquitoes tested | Homozygote mutation (RR) | Heterozygote mutation (RS) | Homozygote wild type (SS) | L1014F *kdr* allele frequency | | $\chi^2$ Test | *P* value |
| --- | --- | --- | --- | --- | --- | --- | --- | --- |
|  |  |  |  |  | *R* | *S* |  |  |
| Fandie | 38 | 38 | 0 | 0 | 1.0 | 0 | - | - |
| Madinagbe | 27 | 9 | 3 | 15 | 0.39 | 0.61 | 15.85 | <0.0001 |
| Maferinyah Centre I | 70 | 69 | 0 | 1 | 0.99 | 0.01 | 70.00 | <0.0001 |
| Moribayah | 6 | 6 | 0 | 0 | 1.0 | 0 | - | - |
| Seneguelen | 46 | 43 | 0 | 3 | 0.93 | 0.07 | 46.00 | <0.0001 |
| Yindi | 55 | 35 | 8 | 12 | 0.71 | 0.29 | 23.05 | <0.0001 |

**Table 5.** G119S *Ace-1* allele frequencies in *An. gambiae* s.l. from six study sites in Forecariah Prefecture, Guinea.

| Study Site | # Mosquitoes tested | Homozygote mutation (RR) | Heterozygote mutation (RS) | Homozygote wild type (SS) | G119S *Ace-1* allele frequency | | $\chi^2$ Test | *P* value |
| --- | --- | --- | --- | --- | --- | --- | --- | --- |
|  |  |  |  |  | *R* | *S* |  |  |
| Fandie | 16 | 0 | 1 | 15 | 0.03 | 0.97 | 0.02 | 0.888 |
| Madinagbe | 4 | 0 | 1 | 3 | 0.13 | 0.88 | 0.08 | 0.777 |
| Maferinyah Centre I | 28 | 0 | 3 | 25 | 0.05 | 0.95 | 0.09 | 0.764 |
| Moribayah | 2 | 0 | 2 | 0 | 0.5 | 0.5 | 2.00 | 0.157 |
| Seneguelen | 9 | 0 | 0 | 9 | 0.0 | 1.0 | - | - |
| Yindi | 8 | 0 | 0 | 8 | 0.0 | 1.0 | - | - |

### Plasmodium falciparum infection and infectivity

The abdomens and head/thoraxes of 484 *An. gambiae* s.l. were screened separately to detect the presence of *Plasmodium falciparum* oocysts and sporozoites, respectively. Overall oocyst and sporozoite rates were 6.0% (29/484) and 0.2% (1/484), respectively. There was no significant difference in oocyst rate among study villages (χ^2^=7.71; *p*=0.173) or molecular forms (χ^2^=0.25; *p*=0.88). In mosquitoes which survived insecticide exposure, oocyst rate was 6.8% (23/340) compared to 4.2% (6/144) among their susceptible counterparts, respectively. The single sporozoite positive individual was collected from Maferinyah Centre I and died following 2X alpha-cypermethrin exposure.

Considering the interaction between pyrethroid resistance and *P. falciparum* infection, there was no significant relationship between oocyst rate and survival following insecticide exposure in all pooled data (χ^2^=0.26; *p*=0.61) or per insecticide (alpha-cypermethrin: χ^2^=0.16; *p*=0.69; deltamethrin: χ^2^=0.07; *p*=0.79 and permethrin: χ^2^=2.28; *p*=0.13). Considering the interaction between carbamate resistance and *P. falciparum* infection, survivors of bendiocarb exposure were significantly more likely to be infected with oocysts (χ^2^=4.86; *p*=0.03).

No significant associations were observed between presence of resistance mutations and oocyst positivity (χ^2^=0.67, *p*=0.72; χ^2^=0.35, *p*=0.56, χ^2^=2.14; *p*=0.34 and χ^2^=0.27; *p*=0.60, for N1575Y, I1527T, L1014F *kdr* and G119S *Ace-1*, respectively).

## Discussion

By 2016, resistance to at least one insecticide has been reported from over 80% of malaria endemic countries^1^, representing a significant threat to the continued efficacy of key malaria control strategies. However, the relative impact of decreased mosquito susceptibility on vectorial capacity remains unknown, largely due to a paucity of field data. In an area of high malaria transmission in Guinea, we characterized levels of insecticide resistance and age of local vector populations, in combination with molecular identification of resistance markers and detection of malaria infection, to begin to address this deficit.

In Forecariah Prefecture, intense pyrethroid resistance was abundant, evidenced by vector populations which were not only resistant to ten times the insecticide concentration required to kill susceptible individuals, but were also capable of surviving these doses for up to two hours. These observations are of significant concern given the coverage of pyrethroid LLINs has been scaled-up in this area. The restoration of mosquito susceptibility following pre-exposure to PBO and the association of L1014F *kdr* with levels of mortality to deltamethrin and permethrin suggests that both target site mutations and over-expression of P450 monooxygenases are contributing to pyrethroid resistance. By comparison, N1575Y and I1527T allele frequencies were lower and neither were associated with increased survival following pyrethroid treatment. This finding aligns with reports from Burkina Faso where upregulated detoxification enzymes were responsible for extreme pyrethroid resistance in *An. coluzzii*, with N1575Y associated with more limited tolerance to deltamethrin^28^. Other studies have proposed that N1575Y may compensate for fitness costs incurred by the L1014F *kdr* mutation and provide additional pyrethroid resistance^29^. The I1527T mutation is located within the III S6 helix of the voltage-gated sodium channel gene, adjacent to a predicted pyrethroid/DDT binding site; nearby residues have already been implicated in resistance in other medically-important vector species^30,31^. However, to date, a role for I1527T in phenotypic resistance in *An. gambiae* s.l. has not been confirmed. While not currently under selection in our study area, future surveillance efforts may wish to monitor changing frequencies of N1575Y and I1527T, in lieu of L1014F *kdr*, which is closely approaching fixation in the majority of our populations, to assess the effect vector interventions are having on the evolution of local, contemporary pyrethroid resistance.

Of the three pyrethroid insecticides under evaluation, resistance levels were highest to permethrin, despite all mass net campaigns in the country having exclusively distributed deltamethrin-treated products^24^. This may be explained by L1014F *kdr* playing a larger contributing role to resistance to type I (permethrin) versus type II (alpha-cypermethrin and deltamethrin) pyrethroids^32^. Levels of bendiocarb resistance were comparatively lower, and not unexpected considering IRS is not routinely conducted by the Guinean NMCP. In the case of carbamate resistance, the presence of the G119S *Ace-1* mutation was highly predictive of bioassay survivorship and tolerance to increased exposure times. Previous studies have demonstrated that the G119S substitution imposes a high fitness cost^33 34^by decreasing affinity of the resistant enzyme for its substrate by more than 60%^35^; heterogeneous or homogeneous duplication of this locus has been proposed to restore activity^36^. In Forecariah Prefecture, frequency of the G119S *Ace-1* resistant allele (of undetermined copy number) was low and likely a result of selective pressure imposed by unregulated agricultural use of carbamates and organophosphates, which were readily available at the local market (S. Irish, personal communication).

Our results demonstrated that pyrethroid resistance was not associated with lower malaria prevalence rates in mosquitoes, with no significant differences observed between rates of *Plasmodium* infection among susceptible and resistant individuals. Our findings contrast with a number of laboratory studies ascribing potential fitness costs to vector insecticide resistance which may have the collateral benefit of reducing malaria transmission^15 17 18 19 20 21^. More worryingly, in our study, survivors of bendiocarb exposure were significantly more likely to be infected with *Plasmodium* oocysts. Pyrethroid exposure has previously been shown to adversely affect *P. falciparum* development in L1014F *kdr* resistant *An. gambiae* s.s. in Uganda^20^, which may explain part of our data, if we assume resistant individuals are frequently surviving contact with LLINs and susceptible vectors are not. Alternative observations have also been reported; *kdr* mutation has been shown to potentiate the vector competence of *An. gambiae* s.s.^22^*, Plasmodium* infection has partially restored the susceptibility of *An. gambiae* carrying the *kdr* mutation to DDT^37^but also reduced the survival of resistant vectors in the absence of insecticide exposure^16^. The discrepancies between *in vitro* studies and our field data may reflect more generalised fitness variation between laboratory mosquito strains and wild populations and/or different underlying resistance mechanisms, e.g. target site mutations *vs*. over-expression of metabolic enzymes, and strongly support the need for additional studies in areas of differing resistance and disease transmission intensities.

We chose to examine the impact of resistance on mosquito age as increasing age has been proposed to restore insecticide susceptibility^38 39 40 41 42 43 44^ and insecticide exposure may still reduce resistant vector life-span through delayed mortality effects^15^. In Forecariah Prefecture, resistant mosquitoes had significantly lower parity rates than their susceptible counterparts. However, a small proportion of those displaying intense resistance (i.e. were capable of surviving exposure to five and ten times the diagnostic dose of pyrethroids) were more likely to be parous. Without additional vector life history information, it is impossible to distinguish between correlation and causality in this scenario. It is tempting to speculate that this small proportion of highly-resistant mosquitoes either evaded or were capable or surviving insecticide exposure for long enough to lay an egg batch. Alternatively, this population could have consisted of more moderately-resistant vectors which underwent multiple sub-lethal contact events, in turn conflating their levels of pyrethroid resistance intensity ^45^.

Elucidating the interaction between insecticide resistance and vectorial capacity is complex and challenging in field conditions and a number of limitations were encountered during this study. Adult female mosquitoes were sampled using human landing catches (HLCs) and via manual aspiration from house walls to maximise the number of individuals available for bioassay testing. However, these strategies may have introduced a bias in species composition collected; previous studies have suggested that proportions of *An. gambiae* s.s. and *An. coluzzii* may differ significantly between larval and adult spray catches^46^. We also encountered issues with individuals that did not amplify with two sub-species PCR assays^47,48^, potentially reflecting polymorphisms at the primer binding sites and/or the presence of a cryptic sub-species; previously a cryptic subgroup GOUNDRY has been reported, with close genetic affinities to *An. coluzzii* and enhanced susceptibility to *P. falciparum* infection49–51. While it is recommended to use F1 progeny mosquitoes for resistance testing^52 53^, to investigate the impact of resistance on vector *Plasmodium* infection, it was necessary to sample wild caught adults of undetermined, mixed age and physiological status (i.e. unfed, fed, gravid etc.); the latter parameter may have varied between sampling method, with more fed mosquitoes collected from inside houses, compared to unfed, host-seeking mosquitoes in HLCs. As a consequence, mortality in our study may have been under-estimated, considering the proposed inverse relationship between vector resistance and age^38 39 40 41 42 43 44^. Similarly, blood-feeding among resistant mosquitoes has been suggested to increase insecticide tolerance^54^, however, it should be noted that no association between physiological status and resistance was observed in our dataset. Our study would have also benefitted from larger sample sizes of mosquitoes tested in bioassays, to statistically power comparisons between different resistance intensities. Furthermore, oocyst positive mosquitoes cannot all be assumed to become infective sporozoite-transmitters, particularly if vector life-span is reduced; it is important to note that a positive abdomen can also reflect a recent feed on an infected individual. Finally, manual parity dissections have a number of known constraints, including insensitivity, especially in low endemicity areas, and inter-operator subjectivity. While every effort was made to consistently bisect individuals, and our reported sporozoite rate was low, other studies have demonstrated higher number of sporozoite false positives by PCR when abdomens were removed posterior to the junction of the abdomen and thorax^55^.

## Conclusions

Study findings present a comprehensive overview of the current levels of insecticide resistance and underlying target site mutations present in Maferinyah, Guinea, an area of high malaria transmission. We describe a methodology to unravel the interaction between insecticide resistance and malaria transmission dynamics, with putative implications for the operational effectiveness of vector control interventions. Local mosquito populations were intensely resistant to pyrethroids (alpha-cypermethrin, deltamethrin and permethrin), associated with high frequencies of the L1014F *kdr* allele. N1575Y and I1527T mutations in the voltage-gated sodium channel gene were present at lower levels and may warrant increased surveillance efforts, particularly as L1014F *kdr* approaches fixation. Restoration of mosquito susceptibility following pre-exposure to PBO indicates upregulated detoxification enzymes are also responsible for extreme pyrethroid resistance in this area and require additional characterization. Despite no ongoing vector control activities using carbamates, bendiocarb resistance was also detected, mediated by the G119S *Ace-1* mutation in a subset of tolerant individuals. Malaria infection (oocyst rate) was not associated with pyrethroid resistance, potentially attributable to the influence chemical exposure may have on parasite development. In general, resistant vectors were younger than their susceptible counterparts; however, a small proportion of intensely resistant mosquitoes were older, which may be cause for concern. Further investigations are necessary to investigate the impact of insecticide resistance on vector fitness, including mosquito fecundity, egg viability, hatchability and parasite development following an infected blood meal.

## Methods

### Study area and mosquito collections

Mosquito collections were undertaken in six villages in the Maferinyah sub-prefecture, Forecariah Prefecture (Fandie, Madinagbe, Maferinyah Centre I, Moribayah, Senguelen and Yindi), in South-West Guinea. Deltamethrin treated bednets were distributed as part of a national mass campaign in Maferinyah in 2016. Sampling was conducted between 22^nd^ June and 17^th^ July 2017, coinciding with the beginning of the long rainy season. Following consent from the household owner, indoor resting, female *Anopheles* mosquitoes were collected from house walls by manual aspiration between 7:00h and 12:00h. In the same villages, mosquitoes were also sampled using human landing catches, carried out between 22:00h and 3:00h. Mosquitoes were stored in cages with access to 10% sugar solution, prior to transport to the Centre de Formation et de Recherche en Santé rurale de Maferinyah (CNFRSR) for analysis.

### CDC resistance intensity and synergist bioassays

All bioassays were performed using mosquitoes identified morphologically as *An. gambiae* s.l. ^56^; wild caught females were held for a maximum of 48 hours before testing. Centers for Disease Control and Prevention (CDC) resistance intensity bioassays for three pyrethroid insecticides (alpha-cypermethrin, deltamethrin and permethrin) were conducted according to published guidelines ^52^. Stock solutions of 1, 2, 5 and 10 times the diagnostic dose of insecticide (alpha-cypermethrin: 12.5 μg/bottle; deltamethrin: 12.5 μg/bottle; and permethrin: 21.5 μg/bottle), were prepared by diluting technical grade insecticide in 50 ml of acetone. Bioassays for bendiocarb were conducted using the diagnostic dose (1X: 12.5 μg/bottle). The inside of each Wheaton 250 ml bottle along with its cap was coated with 1 ml of stock solution by rolling and inverting the bottles. In each test, a control bottle was coated with 1 ml of acetone. Following coating, bottles were left to dry in a dark box. Approximately 15-25 field-caught adult female *An. gambiae* s.l. of unknown age and mixed physiological status, were introduced into each bottle using a mouth aspirator and mortality was recorded at 15 min intervals until all were dead or up to two hours. In select sites with significant pyrethroid resistance, synergist assays were also conducted by pre-exposing mosquitoes to piperonyl butoxide (PBO) for 1 hour (100 μg/bottle) prior to performing bioassays. Multiple replicates were performed per insecticide and study village, depending on mosquito availability, and individual surviving (resistant) and dead (susceptible) mosquitoes were preserved in RNAlater^®^ (Thermo Fisher Scientific, UK) at −20°C at the London School of Hygiene and Tropical Medicine (LSHTM) and CDC. Prior to molecular analysis, mosquito head/thoraxes were separated from abdomens under a dissecting microscope and stored separately.

### Parity dissection

Ovarian dissection to determine mosquito parity was performed on ten mosquitoes, selected randomly from each bottle after bioassay completion ^57^. The ovaries of each mosquito were dissected on a sterile microscope slide in distilled water, using a binocular dissection microscope and physiological status was determined. The ovaries were then examined under a light microscope (10X magnification) for the presence of tightly coiled skeins or loose coils, indicative of a nulliparous or parous ovary, respectively. On completion of ovarian dissection, head/thoraxes and abdomens of each mosquito were stored separately in RNAlater^®^ (Thermo Fisher Scientific, UK) at −20°C.

### Molecular species identification

A subset of susceptible and resistant *An. gambiae* s.l. mosquitoes from all six villages containing both nulliparous and parous individuals were selected for molecular analysis at the LSHTM and CDC. Genomic DNA from dissected body parts was extracted per protocol using Qiagen DNeasy 96 Blood and Tissue kits (Qiagen, UK) at LSHTM or Extracta™ DNA Prep for PCR-Tissue kits (QuantaBio, USA) at CDC.

At LSHTM, molecular species identification was performed using a multiplex TaqMan real time PCR assay to detect and discriminate *An. gambiae* and *An. arabiensis* ^58^. PCR reactions were prepared using Qiagen Quantitect Probes Master mix (Qiagen) with each reaction containing 6.25μl of master mix, a final concentration of 0.8μM of each primer, 0.2μM of probe *An. arabiensis* (Cy5), 80nM of probe *An. gambiae* (FAM) and 1μl of template DNA for a a final reaction volume of 12.5μl. Prepared reactions were run on a Stratagene Mx30005P QPCR system for 10 min at 95°C, followed by 40 cycles of 95°C for 25 sec and 66°C for 60 sec. The increases in the species-specific FAM and Cy5 fluorophores was detected in real-time at the end of each cycle and results were analysed using Stratagene MxPro QPCR software. Positive controls from gDNA extracted from known *An. gambiae* and *An. arabiensis* individuals were included in each run, in addition to no template controls (NTCs). Sub-samples of individuals from each district confirmed as *An. gambiae* were further distinguished as *An. coluzzii* or *An. gambiae* s.s by targeting a *SINE200* insertion only present in *An. coluzzii*^47^. PCR reactions were prepared using Hot Start *Taq* 2X Master Mix (New England Biolabs, UK) with each reaction containing 12.5μl of master mix, a final concentration of 1μM of each primer, 2μl template DNA for a final reaction volume of 25μl. Prepared reactions were amplified using a BIORAD T100 Thermal Cycler for 10 min at 95°C, followed by 35 cycles of 94°C for 30 sec, 54°C for 30 sec, 72°C for 1 min and a final extension of 72°C for 10 min. PCR products were separated and visualised using 2% Egel EX agarose gels (Invitrogen, UK) with SYBR safe and an Invitrogen E-gel iBase Real-Time Transilluminator. *An. coluzzii* individuals with the insertion resulted in a single PCR product of 479 bp and *An. gambiae* s.s. a PCR product of 249 bp.

At CDC, *An. coluzzii* and *An. gambiae* s.s. specimens were differentiated by targeting single nucleotide polymorphisms (SNPs) present in rDNA^48^. PCR reactions were prepared using 20-40ng of DNA, 5X Green GoTaq^®^ Reaction Buffer (Promega, USA), 25mM MgCl_2_, 2.5mM of each dNTP, 1U GoTaq^®^ DNA polymerase and 25 pmol/μl of primers IMP-UN, AR-3T, GA-3T, IMP-S1 and IMP-M1 in a final volume of 25μl. PCR reaction conditions were 95°C for 5 min, followed by 30 amplification cycles (95°C for 30 sec, 58°C for 30 sec, 72°C for 30 sec) and a final elongation step at 72°C for 5 min. Amplified PCR products were visualized on 1.5% agarose gels, stained with GelRed™ (Biotium, USA). *An. arabiensis* Dongola, *An. coluzzii* AKDR and *An. gambiae* s.s. RSP-ST strains from the Malaria Research and Reference Reagent Resource Center (MR4), were used as positive controls. Amplification products of 463bp and 333bp or 463bp and 221bp were indicative of *An. coluzzii* or *An. gambiae* s.s., respectively.

### Insecticide target site mutation detection

Detection of the L1014F West African *kdr* mutation was performed on a subset of individuals from all six districts according to the adapted protocol for allele-specific PCR developed by Martinez-Torres *et al.*^59^. At both LSHTM and CDC, PCR reactions were prepared using Hot Start *Taq* 2X Master Mix (New England Biolabs) with each reaction containing 12.5 μl of master mix and variable final concentrations of primers (IPCF 0.1μM, AltRev 0.1μM, West WT 1μM, West West 1.1μM) for a final reaction volume of 25μl. Prepared reactions were run on a BIO-RAD QPCR system for 5 min at 95°C, followed by 35 cycles of 95°C for 30 sec, 59°C for 30 sec and 72°C for 30 sec and a final extension of 72°C for 5 min. PCR products were separated and visualised using 2% Egel EX agarose gels (Invitrogen) with SYBR safe and an Invitrogen E-gel iBase Real-Time Transilluminator. A PCR product of 214 bp indicated the susceptible wild type allele and a PCR product of 156 bp indicated the resistant allele. *An. coluzzii* AKDR and *An. gambiae* s.s. RSP-ST were used as positive and negative controls for L1014F *kdr*, respectively.

At LSHTM, a larger sub-sample of individuals from all six districts was chosen to be screened for the N1575Y mutation (previously shown to have low prevalence in West African populations) using the TaqMan real time PCR assay developed by Jones *et al*.^29^. PCR reactions were prepared using Qiagen Quantitect Probes Master mix (Qiagen) with each reaction containing 10μl of master mix, a final concentration of 1μM of each primer and 0.5μM of each probe, 5μl of PCR grade water and 2μl of template DNA, for a final reaction volume of 20μl. Prepared reactions were run on a Roche LightCycler 96 System for 15 min at 95°C, followed by 35 cycles of 94°C for 15 sec and 60°C for 60 sec. Positive controls from gDNA extracted from known *An. gambiae s.s*. with or without the N1575 mutation were included on each run in addition to no template controls (NTCs). PCR results were analysed using the LightCycler 96 software (Roche Diagnostics).

At CDC, to detect the N1575Y mutation, a 218 bp fragment of the VGSC channel spanning domains III-IV was sequenced. PCR reactions were prepared using 25μl of 2X AccuStart^™^ II PCR SuperMix, a final concentration of 20pmol/μl of primers Exon_29_F (5’-AAATGCTCAGGTCGGTAAACA-3’) and Exon_29_R (5’-GCCACTGGAAAGAATGGAAA-3’) ^29^ and 2μl of template DNA, for a final reaction volume of 50μl. PCR reaction conditions were 95°C for 3 min, followed by 35 amplification cycles (95°C for 30 sec, 58°C for 30 sec, 72°C for 30 sec) and a final elongation step at 72°C for 5 min. Amplified PCR products were visualized on 2% agarose gels, stained with GelRed™ (Biotium, USA) and purified using the 96-well Millipore™ MultiScreen™ HTS vacuum manifold system. Bi-directional sequencing was performed using primers Exon_29_F and Exon_29_R with the BigDye^®^ Terminator v3.1 Cycle Sequencing Kit (Applied Biosystems, USA) according to the manufacturer’s protocol. Sequencing reactions were purified using the Big Dye^®^ Xterminator™ Purification Kit (Applied Biosystems, USA), according to the manufacturer’s protocol and data were generated using a 3500xL Genetic Analyzer (Applied Biosystems, USA). Sequences were assembled manually in BioEdit v7.0.9.0 sequence alignment editor software (Ibis Biosciences, USA) and unambiguous consensus sequences were produced for each individual. Consensus sequences are available from GenBank under the accession numbers MH929325 - MH929433.

The presence of the *Ace-1* mutation was determined using PCR restriction fragment length polymorphism analysis ^60^. At both LSHTM and CDC, PCR reactions were prepared using Hot Start *Taq* 2X Master Mix (New England Biolabs) with each reaction containing 12.5μl of master mix, a final concentration of 1μM of each primer, 2μl template DNA, to a final reaction volume of 25μl. Prepared reactions were run on a BIO-RAD QPCR system for 5 min at 95°C, followed by 30 cycles of 95°C for 30 sec, 52°C for 30 sec and 72°C for 1 min and a final extension of 72°C for 5 min. PCR products were digested using the *Alu*I restriction enzyme through incubation at 37°C for 16 hr followed by 65°C for 20 min. DNA fragments were visualised on 2% Egel EX agarose gels (Invitrogen) with SYBR safe and an Invitrogen E-gel iBase Real-Time Transilluminator. 194 bp undigested PCR products indicated the susceptible allele and 74 bp and 120 bp digested fragments indicated the presence of the resistant allele. Presence of all three product sizes indicated that the sample was heterozygote.

### Plasmodium falciparum detection

Bisected mosquito head/thoraxes and abdomens were screened separately for *P. falciparum* infection, by targeting a 120 bp sequence of the *P. falciparum* cytochrome c oxidase subunit 1 (*cox1*) mitochondrial gene^61^. At LSHTM real-time PCR reactions were prepared using FastStart SYBR Green Master mix (Roche Diagnostics) with each reaction containing 5μl of master mix, a final concentration of 1μM of each primer, 1μl of PCR grade water and 2μl template DNA, to a final reaction volume of 10μl. Prepared reactions were run on a Roche LightCycler 96 System for 15 min at 95°C, followed by 35 cycles of 95°C for 15 sec and 58°C for 30 sec. Amplification was followed by a dissociation curve (95°C for 10 sec, 65°C for 60 sec and 97°C for 1 sec) to ensure the correct target sequence was amplified. Positive controls from gDNA extracted from a cultured *P. falciparum*-infected blood sample (parasitaemia of ~10%) were included on each run, in addition to no template controls (NTCs). PCR results were analysed using the LightCycler^®^ 96 software (Roche Diagnostics). At CDC, a conventional PCR was used for *cox1* detection^62^. PCR reactions were prepared using 6.25μl of 2X AccuStart™ II PCR SuperMix, a final concentration of 5μM of primers COX-IF (5’-AGAACGAACGCTTTTA ACGCCTG-3’) and COX-IR (5’-ACWGGATGGACTTTATATCCACCATTAAGT-3’) ^62^ and 2μl of template DNA, for a final reaction volume of 12.5μl. PCR reaction conditions were 94°C for 5 min, followed by 40 amplification cycles (94°C for 1 min, 62°C for 1 min, 72°C for 1.5 min) and a final elongation step at 72°C for 10 min. Amplified PCR products were visualized on 2% agarose gels, stained with GelRed™ (Biotium, USA), alongside the same positive and negative controls used in the real-time assay. A positive head/thorax or abdomen was assumed to indicate the presence of sporozoites or oocysts, respectively.

### Data analysis

For all bioassays, mortality was corrected using Abbott’s formula when mortality in the control group was greater than 5% but less than 20%; test results were discarded when control mortality was greater than 20%^63^. Percentage mosquito mortality for each insecticide dose was interpreted using the updated WHO criteria: 98-100% mortality at 30 minutes of exposure indicates ‘susceptibility’, 90-97% mortality suggests ‘possible resistance’ and <90% indicates the presence of ‘resistance’^53^. Pearson’s Chi squared tests and Fisher’s exact tests (when sample sizes were small) were used to investigate the statistical association between various biological variables; the former was also used to detect deviations from Hardy-Weinberg equilibrium. Outcomes from pyrethroid bioassays were evaluated separately to carbamate assays, unless otherwise specified. All statistical analyses were performed in Stata/IC 15.0 (Stata Corp., College Station, USA) with the level of significance set at α=0.05.

### Ethical approval and consent to participate

The study protocol was reviewed and approved by the Comite National d’Ethique pour la Recherche en Sante (030/CNERS/17) and the institutional review boards (IRB) of the London School of Hygiene and Tropical Medicine (#13612 and #14076) and the Centers for Disease Control and Prevention, USA (2018-086). Prior to study initiation, community consent was sought from village leaders and written, informed consent was obtained from the heads of all households selected for participation. Study information was provided to participants in French, *Susu*, *Foula* and *Malinké.* Fieldworkers participating in human landing catches were provided with doxycycline malaria prophylaxis for the duration of the study.

## Acknowledgements

The authors express their sincere thanks to the entomology fieldworkers and the residents of Forecariah Prefecture for their study participation; to Luc Djogbenou and Corine Ngufor for providing G119S *Ace-1* controls; to Martin Donnelly for providing N1575Y controls; to Dustin Miller for providing *An. arabiensis*, *An. coluzzii* and *An. gambiae* s.s. controls; to Alice Sutcliffe for providing primers and reagents; and to Jo Lines and Cheryl Whitehorn for providing training and support. Study funding was provided by a Royal Society of Tropical Medicine and Hygiene small grant, a Sir Halley Stewart Trust grant, a L’Oreal-UNESCO for Women in Science UK and Ireland Fellowship awarded to LAM, individual Bayer Vector Control Research and Travel grants awarded to EC and NMV, the Helena Vrbova scholarship awarded to NMV and a Wellcome Trust /Royal Society grant awarded to TW (101285/Z/13/Z): http://www.wellcome.ac.uk; https://royalsociety.org.

## Author Contributions

LAM, TW, MS and AHB designed the study and were responsible for data analysis and interpretation. EC, NMV and MS led the entomology field activities and participated in data collection. EC, NVM, TW, JO and LAM performed the molecular assays. LAM, TW, EC and NMV drafted the manuscript which was revised by co-authors. All authors read and approved the final manuscript.

